# Non-canonical activation of IRE1α during *Candida albicans* infection enhances macrophage fungicidal activity

**DOI:** 10.1101/2023.10.02.560560

**Authors:** Michael J. McFadden, Mack B. Reynolds, Britton C Michmerhuizen, Einar B. Ólafsson, Faith M. Anderson, Tracey L. Schultz, Mary X.D. O’Riordan, Teresa R. O’Meara

**Affiliations:** Department of Microbiology and Immunology, University of Michigan, Ann Arbor, MI 48109, USA

## Abstract

While the canonical function of IRE1α is to detect misfolded proteins and activate the unfolded protein response (UPR) to maintain cellular homeostasis, microbial pathogens can also activate IRE1α, which modulates innate immunity and infection outcomes. However, how infection activates IRE1α and its associated inflammatory functions have not been fully elucidated. Recognition of microbe-associated molecular patterns can activate IRE1α, but it is unclear whether this depends on protein misfolding. Here, we report that a common and deadly fungal pathogen, *Candida albicans*, activates macrophage IRE1α through C-type lectin receptor signaling, reinforcing a role for IRE1α as a central regulator of host responses to infection by a broad range of pathogens. This activation did not depend on protein misfolding in response to *C. albicans* infection. Moreover, lipopolysaccharide treatment was also able to activate IRE1α prior to protein misfolding, suggesting that pathogen-mediated activation of IRE1α occurs through non-canonical mechanisms. During *C. albicans* infection, we observed that IRE1α activity promotes phagolysosomal fusion that supports the fungicidal activity of macrophages. Consequently, macrophages lacking IRE1α activity displayed inefficient phagosome maturation, enabling *C. albicans* to lyse the phagosome, evade fungal killing, and drive aberrant inflammatory cytokine production. Mechanistically, we show that IRE1α activity supports phagosomal calcium flux after phagocytosis of *C. albicans*, which is crucial for phagosome maturation. Importantly, deletion of IRE1α activity decreased the fungicidal activity of phagocytes *in vivo* during systemic *C. albicans* infection. Together, these data provide mechanistic insight for the non-canonical activation of IRE1α during infection, and reveal central roles for IRE1α in macrophage antifungal responses.

## Introduction

Intracellular infection by diverse pathogens triggers cell stress programs, such as the unfolded protein response (UPR), whose three branches (IRE1α, PERK, and ATF6) have broad consequences for host antimicrobial defenses through regulation of innate immunity, cellular metabolism and homeostasis, and cell differentiation or cell death pathways^1–4^. Canonically, accumulation of misfolded proteins in the ER lumen triggers activation of the UPR^5^. As proteostasis is required for cellular function, UPR activation can restore cellular homeostasis by modulating gene expression to promote protein folding and expansion of the ER network. Alternatively, failure to overcome proteotoxic stress leads to cell death^5^. Therefore, initiation of the UPR during infection may be critical to circumvent the effects of pathogen virulence factors, support the production of secreted proteins such as cytokines through cooperation with proinflammatory transcription factors, or to regulate organelle contact sites for inter-organelle communication^4,6,7^. Still, the utility and effects of UPR activation during infection are not fully understood and differ in response to individual pathogens, which may differentially exploit UPR activation for pathogenesis^8–15^.

After detecting misfolded protein accumulation in the ER lumen, IRE1α assembles into small oligomers that allow its *trans*-autophosphorylation^16^. Autophosphorylation of IRE1α results in activation of its endonuclease domain, allowing IRE1α to remove a short intronic sequence from the *Xbp1* transcript in a non-canonical mRNA splicing reaction, orthologous to the Ire1-*Hac1* splicing reaction that drives the UPR in yeast^17–19^. *Xbp1* splicing results in a frameshift within the open reading frame, allowing translation and protein synthesis of the transcription factor XBP1S, which promotes the transcription of genes involved in ER quality control^20^. However, the IRE1α branch of the UPR can also be selectively triggered by infection or detection of microbe-associated molecular patterns (MAMPs) by pattern recognition receptors (PRRs), such as Toll-like receptors (TLRs)^6,21^. Additionally, the regulatory roles of IRE1α extend beyond XBP1S, as IRE1α itself can modulate JNK pathway activation, orchestrate organelle contact sites, and regulate metabolic plasticity^23,26,27,74^. Previous reports suggested that protein misfolding-independent activation of IRE1α may occur following TLR stimulation, although this model has not been directly tested^6,21^.

IRE1α has broad regulatory roles and consequences for infection and immunity. For example, the IRE1α-XBP1S axis can promote the expression of proinflammatory cytokines^6,22^, modulate metabolic plasticity^24^, and promote ER homeostasis^25^ during infection. Additionally, IRE1α can facilitate intra-organelle communication for ER-mitochondria calcium signaling and promotion of reactive oxygen species (ROS) generation^12,13,26^. Through its regulatory effects on gene expression, metabolism, and redox balance, IRE1α can promote bacterial killing or inflammasome activation in phagocytic cells^10,12^. Despite these known roles of IRE1α in bacterial and viral infection, mechanistic understanding of IRE1α activation during infection is lacking. Further, our understanding of the role of IRE1α during fungal infection is only beginning to emerge.

Given its many functions in host responses to infection, we sought to understand the role of IRE1α in macrophage interactions with *Candida albicans*. *C. albicans* is a common fungal member of the human mucosal microbiota and an opportunistic pathogen^28^. Phagocytic cells are an important early line of defense against systemic infection by *C. albicans*^29^. Macrophages and neutrophils can recognize and phagocytose *C. albicans* predominantly through C-type lectin receptor (CLR) signaling and eliminate infection through fungicidal activity or secretion of cytokines to orchestrate antifungal immunity^30–33^. Interestingly, a recent report found that IRE1α can be activated in neutrophils upon *C. albicans* infection, and IRE1α activity contributes to the immunopathology of systemic *C. albicans* infection^14^, revealing the importance of regulation of this pathway during infection. However, the role of IRE1α in macrophage responses to *C. albicans* have not been investigated. Macrophages are crucial for early antifungal responses *in vivo* and are thought to control *C. albicans* dissemination through phagocytosis, direct antifungal activity, and cytokine signaling to recruit neutrophils to sites of infection^29,34,35^. During intracellular growth in macrophages, *C. albicans* hyphal formation can allow it to escape the phagosome and kill macrophages through lysis or programmed cell death through pyroptosis^36–41^. However, the mechanisms by which macrophages contain and kill *C. albicans* are incompletely understood. Indeed, levels of microbicidal effectors, such as ROS, are not reliable predictors of phagocyte fungicidal activity^42,43^. Recent work reported that lysosome fusion with the expanding *C. albicans*-containing phagosome is crucial to maintain phagosome integrity, prevent phagosomal rupture, and allow fungicidal activity^43–46^. Together, these findings suggests that phagosome maturation is a critical component of antifungal responses by macrophages.

Here, we report that IRE1α is activated in macrophages following infection by *C. albicans*. Importantly, IRE1α activation was dependent on CLR signaling, but did not depend on detectable accumulation of misfolded proteins, suggesting a non-canonical mechanism of activation. Additionally, we found IRE1α is dispensable for phagocytosis of *C. albicans* by macrophages, but contributes to their fungicidal activity *in vitro* and *in vivo*. Macrophages lacking IRE1α activity failed to efficiently recruit lysosomes to the phagosome, which was followed by increased phagosome rupture and more hyphal growth by *C. albicans*. These results reveal a role for IRE1α in the fungicidal capacity of macrophages, advancing our understanding of the emerging role of IRE1α in antifungal immunity.

## Results

### *C. albicans* infection results in activation of macrophage IRE1α

While the ER stress sensor IRE1α is activated in response to bacterial and viral infection, its role and activation in response to fungal infections is only beginning to emerge. To determine whether macrophage IRE1α is activated during *C. albicans* infection, we measured splicing of *Xbp1* mRNA in immortalized bone marrow-derived macrophages (iBMDM) infected with *C. albicans*, or treated with known IRE1α activating stimuli, bacterial lipopolysaccharide (LPS) or thapsigargin, as positive controls. Using semi-quantitative RT-PCR analysis of *Xbp1* mRNA, we observed that *C. albicans* infection induces *Xbp1* splicing in wild-type iBMDM (WT), albeit to a lesser extent than the positive controls LPS and thapsigargin (Fig. 1A). *Xbp1* splicing did not occur in response to any of the treatments in a clonal iBMDM cell line lacking exons 20 and 21 of IRE1α (IRE1^ΔR^), which are required for its endonuclease activity^47^ (Fig. 1A, 1B). Analysis of *Xbp1* splicing by RT-qPCR following a timecourse of *C. albicans* infection showed induction of *Xbp1-S* at 4 hours post-infection (hpi) with *C. albicans* (Fig. 1C). As the SC5314 reference strain can be an outlier in virulence and hyphal formation^28,48^, we measured *Xbp1-S* induction following infection with a selection of commensal *C. albicans* isolates previously isolated from healthy donors^28^ and demonstrated that all isolates resulted in comparable *Xbp1* splicing to the reference strain SC5314 (Fig. 1D).

**Figure 1:**
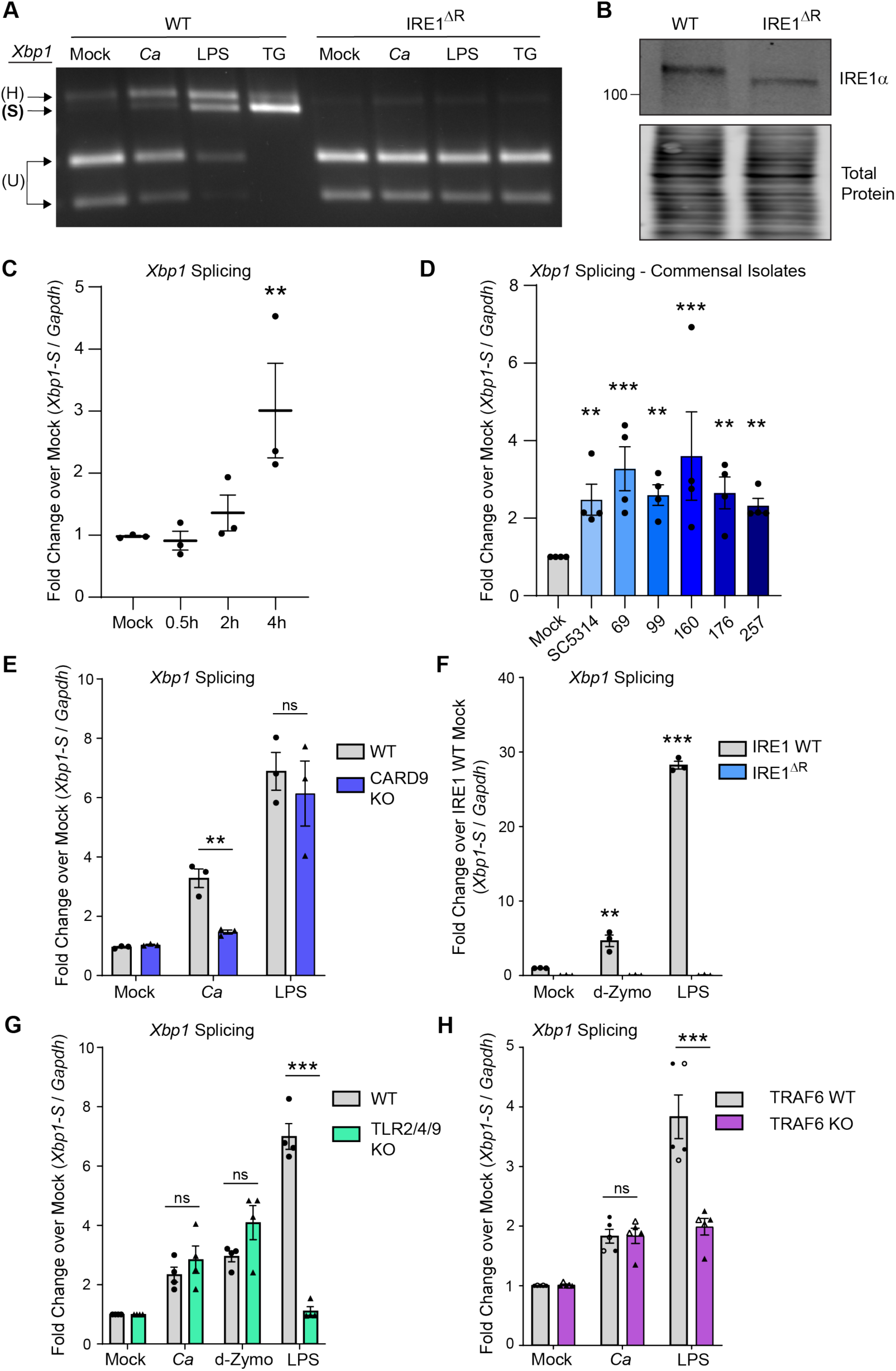
*C. albicans* infection results in activation of macrophage IRE1α. **(A)** iBMDM cell lines (WT or IRE1^ΔR^) were infected with *C. albicans* (MOI=1), treated with LPS (100 ng/mL), or thapsigargin (5 µM), or mock treated for 4 hours. *Xbp1* mRNA splicing was measured by semi-quantitative RT-PCR amplification of the *Xbp1* transcript as a readout of IRE1α activity. **(B)** Immunoblot analysis of lysates from WT or IRE1^ΔR^ iBMDM cell lines to confirm IRE1α truncation in IRE1^ΔR^ cells resulting from removal of floxed exons 20 and 21. **(C)** Expression of the short isoform of *Xbp1* was measured using RT-qPCR over a timecourse following *C. albicans* infection of iBMDM (MOI=1). **(D)** Expression of the short isoform of *Xbp1* at 4 hours post-infection with commensal *C. albicans* isolates as well as the lab strain SC5314 (MOI=1) was measured using RT-qPCR. **(E)** Expression of the short isoform of *Xbp1* was measured using RT-qPCR at 4 hours following *C. albicans* infection (MOI=1) or LPS treatment (100 ng/mL) of WT or CARD9 KO iBMDM. **(F)** Expression of the short isoform of *Xbp1* was measured using RT-qPCR at 4 hours following *C. albicans* infection (MOI=1), depleted Zymosan treatment (d-Zymo; 100 µg/mL) to stimulate Dectin-1, or LPS treatment (100 ng/mL) of WT or IRE1^ΔR^ iBMDM. **(G)** Expression of the short isoform of *Xbp1* was measured using RT-qPCR at 4 hours following *C. albicans* infection (MOI=1), LPS treatment (100 ng/mL), or depleted Zymosan treatment (d-Zymosan; 100 µg/mL) of WT or TLR2/4/9 KO iBMDM. **(H)** Expression of the short isoform of *Xbp1* was measured using RT-qPCR at 4 hours following *C. albicans* infection or LPS treatment of two pairs of clonal iBMDM (WT or TRAF6 KO; MOI=1). Closed symbols are data from WT-1 and KO-1; open symbols are data from WT-2 and KO-2. Data are representative of 3-4 individual experiments. Graphs show the mean ± SEM of biological replicates (C-H). *p < 0.05, **p < 0.01, ***p < 0.005 by 2-way ANOVA of log-transformed data with Sidak’s multiple comparisons test. ns, not significant.

*Xbp1* splicing leads to translation of the transcription factor XBP1S to induce the transcription of ER quality control responsive genes following unfolded protein stress. However, while LPS and thapsigargin treatment led to accumulation of XBP1S by 4 hpi, infection with *C. albicans* did not lead to induction of XBP1S protein expression at 4, 6, or 8 hours post-infection (Fig. S1). Thus, IRE1α function during *C. albicans* infection of macrophages is likely independent of the transcription factor XBP1S. Together, these results indicate that *C. albicans* infection results in mild activation of IRE1α in macrophages.

### C-type lectin receptor signaling drives TRAF6-independent IRE1α activation during *C. albicans* infection

C-type lectin receptors, which detect components of the cell wall of *C. albicans*^31^, are the major pattern recognition receptor for recognition of *C. albicans* in macrophages^49^. To determine whether C-type lectin receptor (CLR) signaling contributes to IRE1α activation during *C. albicans* infection, we measured *Xbp1* splicing in iBMDM lacking the CLR signaling adaptor protein CARD9 (CARD9 KO), compared to WT iBMDM. CARD9 was required for *Xbp1* splicing in response to *C. albicans*, but dispensable for *Xbp1* splicing in response to LPS, which activates a distinct signaling pathway through Toll-like receptor 4 (TLR4)^50,51^ (Fig. 1E). These results suggest CLR signaling is required for IRE1α activation in response to *C. albicans*. Next, we addressed whether CLR agonism is sufficient to stimulate IRE1α activity by treating WT or IRE1^ΔR^ iBMDM with a Dectin-1 specific agonist, depleted Zymosan (d-Zymosan). We found that depleted Zymosan treatment was sufficient to trigger IRE1α-dependent *Xbp1* splicing, demonstrating that CLR agonism triggers IRE1α activity (Fig. 1F). Similar to results with *C. albicans* infection, *Xbp1* processing by IRE1α was more strongly stimulated by LPS than by depleted Zymosan (Fig. 1F). Despite CLRs being the major pattern recognition receptors for *C. albicans,* TLRs can also respond to fungal cells^52^. To test whether TLR engagement is necessary for IRE1α activation in response to *C. albicans*, we measured *Xbp1* splicing in BMDM lacking TLR2, TLR4, and TLR9 (TLR2/4/9 KO). We observed a similar level of *Xbp1* splicing to WT iBMDM in response to *C. albicans* and depleted Zymosan, although *Xbp1* splicing was ablated in response to LPS, as expected (Fig. 1G). Together, these results suggest CLR signaling is necessary and sufficient for IRE1α activation in response to *C. albicans*.

TRAF6 is a crucial E3 ubiquitin ligase involved in innate immune signaling for both TLR and CLR pathways^53,54^. This ubiquitin ligase can directly ubiquitinate IRE1α and facilitates the ubiquitination and activation of IRE1α after LPS treatment^6,21^. Therefore, we tested whether TRAF6 is involved in IRE1α activation in response to *C. albicans* infection. While knockout of TRAF6 resulted in the expected decrease in *Xbp1* splicing in response to LPS, we observed that *Xbp1* splicing in response to *C. albicans* infection was not affected by TRAF6 deletion (Fig. 1H). Therefore, CLR-mediated IRE1α activation is TRAF6-independent. These data reveal that CLR signaling through the adaptor protein CARD9 triggers IRE1α activation independently of TLR signaling or TRAF6, in contrast to LPS-driven IRE1α activation, which depends on TLR signaling to TRAF6. These data reveal a distinct mechanism of IRE1α activation in macrophages through CLR signaling during fungal infection.

**Figure S1: Related to Figure 1.**
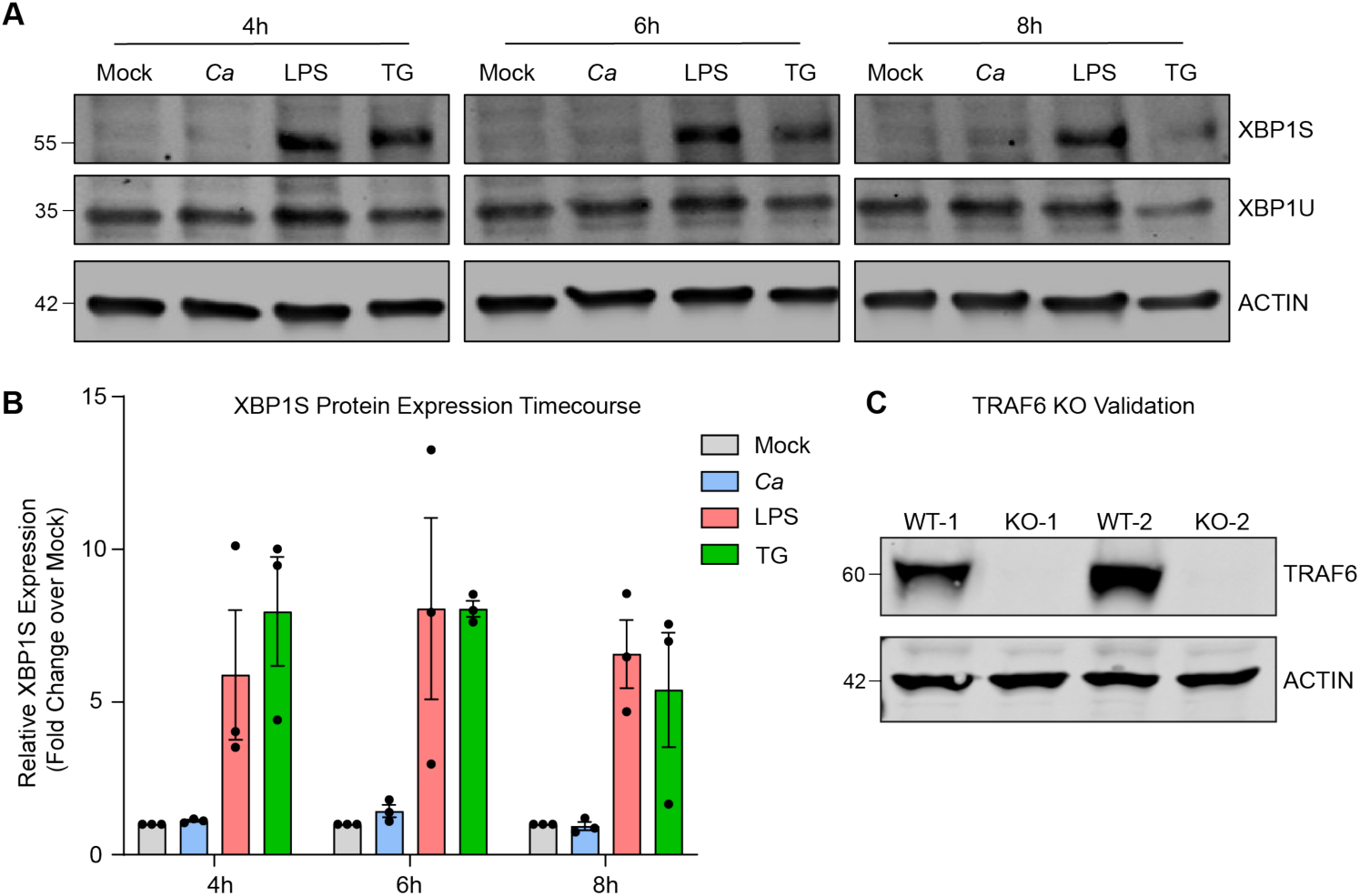
**(A)** Immunoblot analysis of XBP1S and XBP1U expression from WT iBMDM lysates following infection with *C. albicans* (MOI=1), or treatment with positive controls LPS (100 ng/mL) or thapsigargin (5 µM). **(B)** Quantification of 3 independent experiments, as shown in (A). **(C)** Immunoblotting validation of clonal TRAF6 WT controls (WT-1 and WT-2) and TRAF6 knockout iBMDM (KO-1 and KO-2).

### PRR-mediated activation of IRE1α occurs independently of misfolded protein stress

A potential mechanism for CLR-mediated IRE1α activation is by overwhelming protein folding capacity of the ER due to increased cytokine production, leading to protein misfolding and thus IRE1α activation. To test this hypothesis and determine whether new gene synthesis is required for IRE1α activation during *C. albicans* infection, we inhibited transcription or translation during infection with *C. albicans* or during treatment with thapsigargin. Surprisingly, neither inhibition of transcription nor translation, using actinomycin D or cycloheximide treatment, respectively, inhibited *Xbp1* splicing during *C. albicans* infection (Fig. 2A-B). Translation inhibition using cycloheximide was sufficient to alleviate *Xbp1* splicing specifically in response to thapsigargin, likely by reducing the nascent protein folding burden (Fig. 2B). These data indicate that new gene synthesis does not contribute to IRE1α activation during *C. albicans* infection, and presented the intriguing possibility that *C. albicans* infection does not induce unfolded proteins. Indeed, this possibility has been suggested for TLR-driven IRE1α activation, but was not previously directly tested^6,21^. To specifically test this, we measured whether misfolded proteins accumulate during PRR-mediated activation of IRE1α by either *C. albicans* infection or LPS treatment. Thioflavin T (ThT) is widely used to detect protein misfolding, as it exhibits increased fluorescence in the presence of misfolded proteins^55^. While ThT intensity showed an expected increase at 2 hours-post thapsigargin treatment, neither *C. albicans* infection nor LPS treatment increased ThT intensity over mock treatment (Fig. 2C-D). Further, neither *C. albicans* infection nor LPS treatment led to increased ThT intensity at 4 hpi, suggesting IRE1α activation occurs without accumulation of misfolded proteins during these responses (Fig. 2E). Even at 8 hpi, *C. albicans* infection did not induce protein misfolding (Fig. 2F). While LPS treatment did lead to increased protein misfolding at 8 hours post-treatment (Fig. 2F), this occurred after the robust IRE1α activation observed at 4 hours post-treatment (Fig. 1A). Finally, we measured induction of UPR-responsive genes by RT-qPCR in response to *C. albicans* infection, LPS and depleted zymosan treatment, or thapsigargin treatment (Fig. 2G-H). *C. albicans* infection and depleted zymosan treatment did not lead to induction of UPR-responsive genes (*Ddit3*, *Grp78*, *Grp94*, and total *Xbp1*) at 4 or 6 hours. Similarly, LPS treatment did not lead to global induction of UPR-responsive genes, and only led to significant induction of *Grp78* at 4 hours and *Xbp1-T* at 6 hours. Conversely, thapsigargin treatment triggered induction of all of these genes at both 4 and 6 hours post-treatment, as expected (Fig. 2G-H). Together, these data suggest that while protein misfolding can occur in response to microbial stimuli, it is not needed to trigger IRE1α activation during innate immune responses, and points to a non-canonical mode of IRE1α activation during infection.

**Figure 2:**
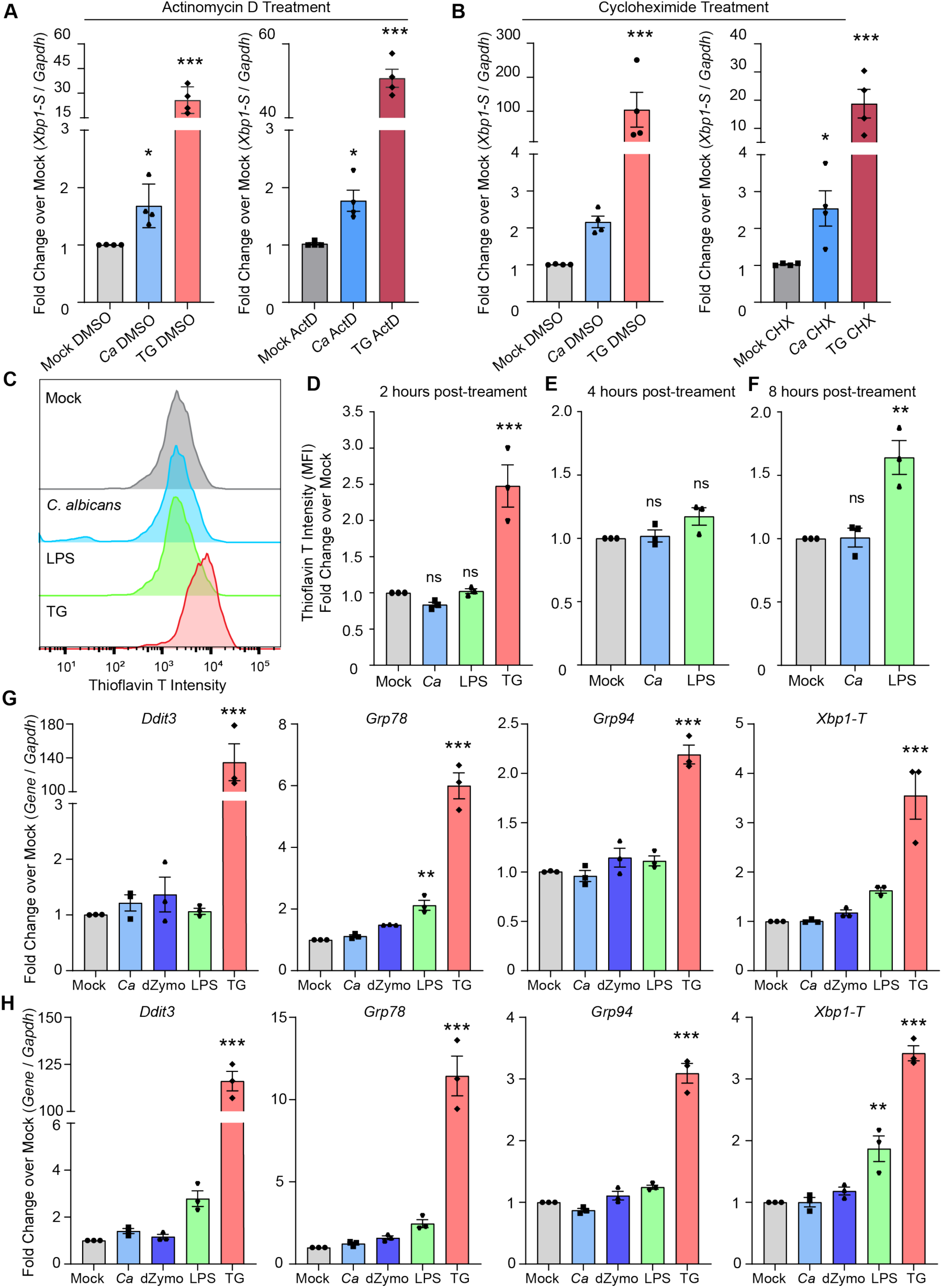
PRR-mediated activation of IRE1α occurs independently of misfolded protein stress. **(A-B)** Expression of the short isoform of *Xbp1* was measured using RT-qPCR following *C. albicans* infection (MOI=1) of iBMDM or treatment with thapsigargin (TG; 5 µM) as a control, compared to mock treatment. Actinomycin D (ActD; 20 µM) was used to inhibit new transcription during treatments (A), and cycloheximide (CHX; 10 µM) was used to inhibit translation during treatments (B), and relative fold changes were measured over matched mock samples. **(C)** Representative graphs showing fluorescence intensity of Thioflavin T (ThT) measured by flow cytometry to quantify protein misfolding in iBMDM following infection by *C. albicans* (MOI=1), or treatment with LPS (100 ng/mL) or thapsigargin (TG; 5 µM) as a positive control. **(D-F)** Quantification of ThT fluorescence intensity at 2 hours (D), 4 hours (E), or 8 hours post-indicated treatment, shown as fold change over mock. **(G-H)** Expression of UPR-responsive genes was measured using RT-qPCR at 4 hours (G) or 6 hours (H) following *C. albicans* infection (MOI=1), depleted Zymosan treatment (d-Zymosan; 100 µg/mL), LPS treatment (100 ng/mL), or thapsigargin treatment (TG; 5 µM). Graphs show the mean ± SEM of biological replicates. *p < 0.05, **p < 0.01, ***p < 0.005 by 2-way ANOVA with Sidak’s multiple comparisons test of log-transformed data (A, B), one-way ANOVA with Tukey’s multiple comparisons test (D-F), or one-way ANOVA with Dunnett’s multiple comparisons test (G-H).

### IRE1α promotes phagosome maturation during C. albicans infection

To explore the potential roles of IRE1a in macrophage antifungal functions, we first tested the ability of IRE1 WT and IRE1^ΔR^ macrophages to ingest *C. albicans* through phagocytosis. Interestingly, IRE1^ΔR^ macrophages showed increased efficiency of *C. albicans* phagocytosis (Fig. 3A-B). Following phagocytosis of large particles, the ER is thought to regulate phagosome maturation through poorly understood mechanisms^56^. Importantly, phagosome maturation is required for containment of *C. albicans* hyphae within the phagosome, as lysosome fusion allows membrane donation to support expansion of the phagosome^44^. Therefore, we next tested whether IRE1^ΔR^ macrophages showed impaired phagosome maturation during *C. albicans* infection by measuring recruitment of the lysosomal protein LAMP1 to the phagosome containing *C. albicans* (Fig. 3C-D). IRE1 WT macrophages recruited LAMP1 to the phagosome by 2 hpi, but IRE1^ΔR^ macrophages infected with *C. albicans* failed to efficiently recruit LAMP1 to the phagosome (Fig. 3C-D). However, overall lysosome biogenesis did not appear to be impaired in IRE1^ΔR^ macrophages, as labeling acidic cellular compartments with LysoSensor Blue/Yellow dye showed IRE1^ΔR^ macrophages had similar acidity as IRE1 WT macrophages, and *C. albicans* infection or ammonium chloride treatment led to the expected alkalinization of both cell lines ^46,57–59^ (Fig. 3E). These data suggest that IRE1α activity is specifically required for efficient phagolysosomal fusion during *C. albicans* infection.

**Figure 3:**
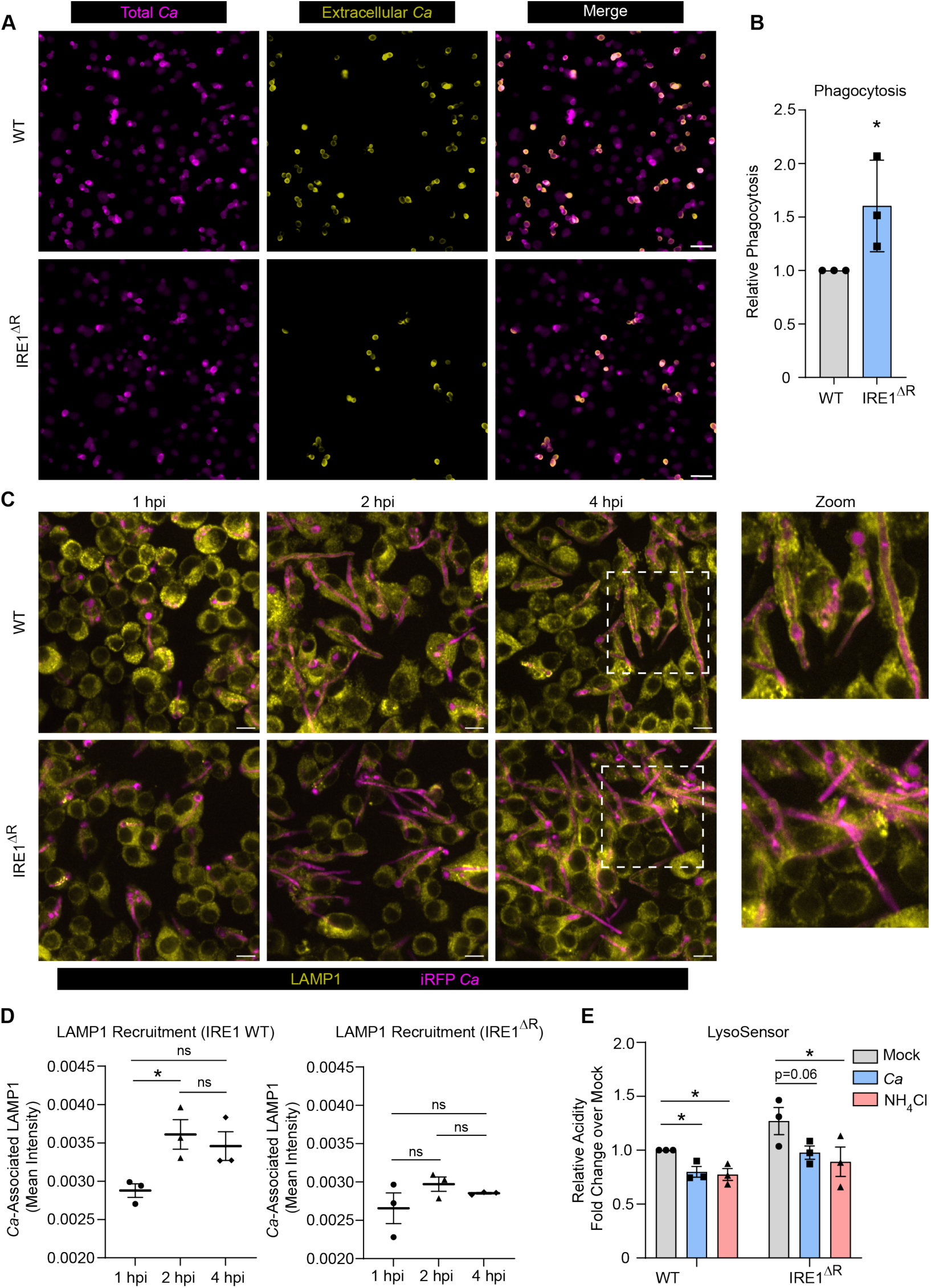
IRE1α promotes phagosome maturation during *C. albicans* infection. **(A)** Representative phagocytosis assay micrographs show total *C. albicans* (intracellular and extracellular; magenta) and extracellular *C. albicans* (yellow) following 30 minutes of phagocytosis by iBMDM (IRE1 WT or IRE1^ΔR^). Scale bar 10 µm. **(B)** Quantification of 3 independent phagocytosis experiments. Relative phagocytosis was calculated as fold change over IRE1 WT. **(C)** Representative images showing LAMP1 (yellow) recruitment to phagosomes containing iRFP-expressing *C. albicans* (magenta) in IRE1 WT or IRE1^ΔR^ iBMDM at indicated times post-infection. **(D)** Quantification of LAMP1 recruitment to phagosomes containing *C. albicans* in IRE1 WT or IRE1^ΔR^ iBMDM, as measured by LAMP1 mean fluorescence intensity associated with *C. albicans*-expressed iRFP. **(E)** The relative acidity of IRE1 WT or IRE1^ΔR^ iBMDM following infection with *C. albicans* or treatment with NH4Cl as a control, shown as the relative ratio of LysoSensor intensity at acidic (Excitation 384 nm, Emission 540 nm) and basic (Excitation 329 nm, Emission 440 nm) conditions. Values are the mean ± SEM from 3 biological replicates. *p < 0.05, **p < 0.01, *** p < 0.001, by unpaired Student’s t-test (B) or one-way ANOVA with Tukey’s multiple comparisons test (D, E).

### IRE1α promotes phagosomal calcium flux necessary for phagosome maturation

To understand of the role of IRE1α in phagosome maturation, we investigated its impact on gene expression during *C. albicans* infection or mock treatment using RNA sequencing. Despite not observing robust XBP1S induction during *C. albicans* infection (Fig. S1), we reasoned that IRE1α activity could modulate gene expression by XBP1S-independent mechanisms, such as cleavage of other transcripts or microRNAs^60–62^, or interaction with ER-localized RNA species under homeostatic or ER stress conditions^63^. As expected, we found many differentially regulated genes in IRE1^ΔR^ macrophages during *C. albicans* infection (Fig. S2A; Table S1.1) compared with WT macrophages at 4 hpi. Importantly, the IRE1^ΔR^ macrophages had broadly similar expression of ER stress-related genes as WT control macrophages, suggesting that lack of IRE1α does not result in chronic ER stress during infection (Fig. S2B). Interestingly, gene ontology analysis revealed that genes involved in endocytosis and calcium homeostasis were enriched among downregulated genes in IRE1^ΔR^ macrophages (Table S1.2). These included genes involved in ER homeostasis (*Kctd17*, *Atp2a3*, *Gramd2*)^64–66^, as well as the major lysosome calcium channels *Mcoln1* and *Mcoln3*^67^ (Fig. S2C). However, the expression of genes involved in general cellular calcium uptake and homeostasis, such as *Calm1*, *Calr*, *Stim1*, *Orai1-3*, and *Ryr1* and *Ryr3* was similar in WT and IRE1^ΔR^ macrophages (Fig. S2C). Therefore, we hypothesized that organellar calcium signaling may be specifically impaired in IRE1^ΔR^ macrophages. Calcium flux regulates phagosome formation and maturation^68^ and is required for lysosome recruitment to the phagosome during *C. albicans* infection^44^. We investigated whether calcium flux is perturbed in IRE1^ΔR^ macrophages during phagocytosis of *C. albicans* using the fluorescent calcium ion indicator Fluo4-AM. In WT macrophages, calcium flux was observed during macrophage-*C. albicans* interactions and during phagocytosis of *C. albicans* (Supplemental Movie 1, Fig. 4A). At baseline, WT and IRE1^ΔR^ macrophages showed similar Fluo4 fluorescence intensity, suggesting calcium stores are not depleted in IRE1^ΔR^ macrophages (Fig. 4B). Additionally, we measured cellular calcium flux per cell in WT and IRE1^ΔR^ macrophages following *C. albicans* infection (Supplemental Figure 3A). Cellular calcium flux was comparable between WT and IRE1^ΔR^ macrophages, as similar frequency of cellular calcium flux was observed (Fig. 4C), as well as similar ‘excitability’ of macrophages during *C. albicans* infection (Fig. 4D). However, shortly after phagocytosis, phagosomal calcium influx was frequently observed in WT macrophages (Supplemental Movie 1, Fig. 4A), seen as a clear but transient ring around the engulfed yeast. However, phagosomal calcium flux was rarely observed in IRE1^ΔR^ macrophages after phagocytosis of *C. albicans* (Supplemental Movie 2, Fig. 4E). Indeed, quantification of phagosomal calcium flux at 20 minutes post-infection in WT macrophages revealed that roughly half of macrophages that had phagocytosed *C. albicans* had active phagosomal calcium flux, with phagosomal intensity above that of the cytosol, whereas less than 20 percent of IRE1^ΔR^ macrophages showed phagosomal calcium flux (Fig. 4F). Additionally, the fluorescence intensity of the *C. albicans* phagosome relative to the cytosol was higher in WT macrophages than in IRE1^ΔR^ macrophages (Fig. 4G). Together, these data suggest that phagosomal calcium flux is specifically impaired in IRE1^ΔR^ macrophages, possibly due to defective expression of calcium signaling-related genes. Previous work showed that phagosome-derived calcium is required for lysosome recruitment during *C. albicans* infection, and that calcium chelation disrupts phagosome maturation^44^. Therefore, to test the hypothesis that calcium flux is required for maturation of *C. albicans*-containing phagosomes, we treated macrophages with a cell permeable calcium chelator, BAPTA-AM, during *C. albicans* infection. BAPTA-AM treatment impaired phagosome maturation in WT macrophages, while defective phagosome maturation observed in IRE1^ΔR^ macrophages was not further impacted by BAPTA-AM treatment (Fig. 4H). These data suggest that defective phagosomal calcium flux in IRE1^ΔR^ macrophages perturbs phagosome maturation.

**Figure 4:**
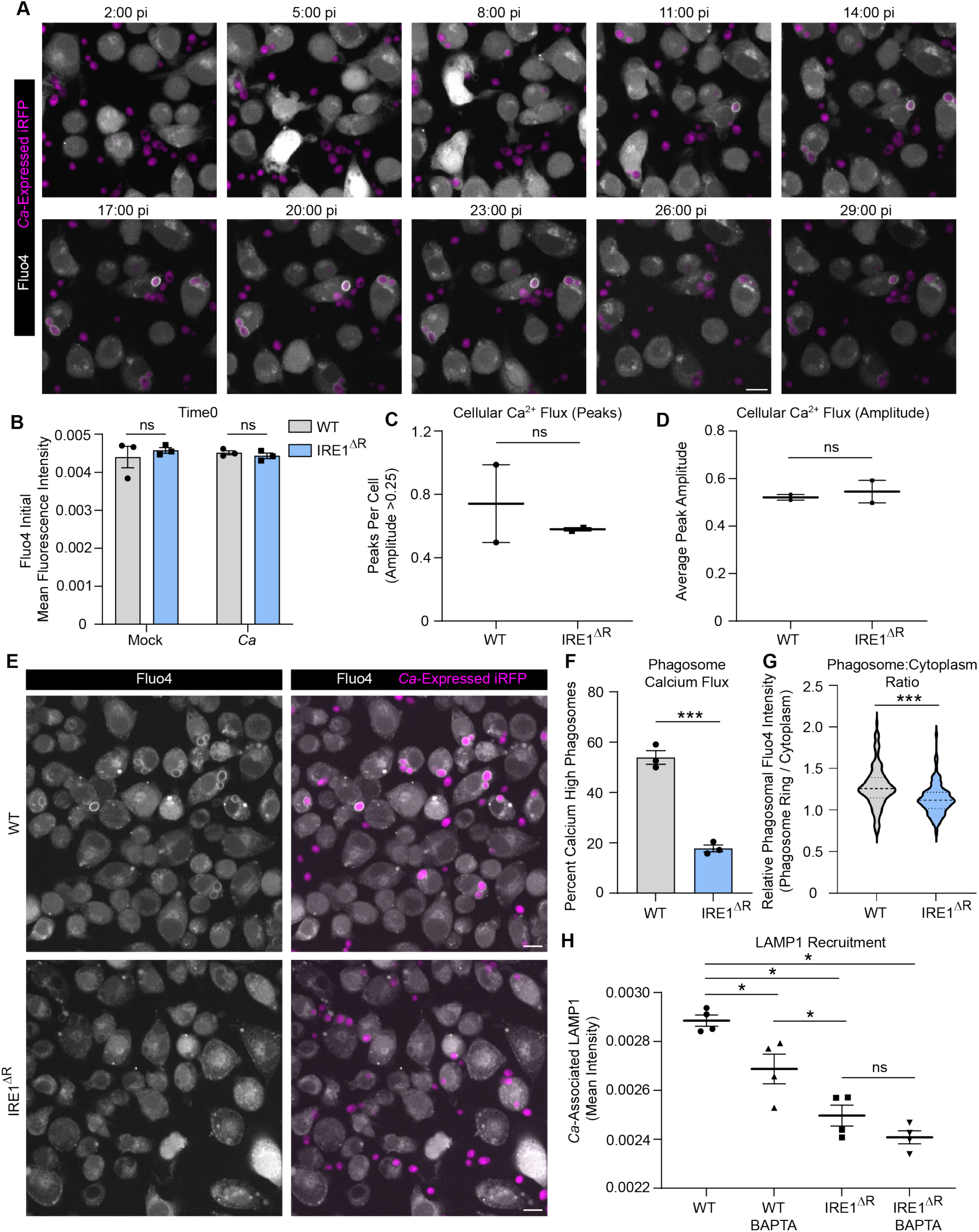
IRE1α promotes phagosomal calcium flux necessary for phagosome maturation. **(A)** Representative micrographs from timelapse imaging of Fluo4 during *C. albicans* infection of WT iBMDM. Scale bar 10 µm. **(B)** Mean fluorescence intensity of Fluo4 at time 0, or the beginning of live imaging, in WT or IRE1^ΔR^ iBMDM. **(C-D)** Analysis of cellular calcium flux of WT or IRE1^ΔR^ iBMDM during early interactions with *C. albicans*. Graphs show the number of peaks per cell, defined as ≥0.25 increase in normalized Fluo4 fluorescence (C), or the average amplitude of peaks (D). **(E)** Representative micrographs of WT or IRE1^ΔR^ iBMDM following phagocytosis of *C. albicans* (20 mins post-infection; MOI 2) showing early cellular calcium flux, and influx of calcium specifically in the phagosome following phagocytosis of *C. albicans*. Scale bar 10 µm. **(F)** Quantification of calcium-high phagosomes, defined by a 1.25-fold increase of the mean fluorescence intensity of the cell (20 mins post-infection; MOI 2). **(G)** Violin plot of the ratio of phagosomal to cytosolic mean fluorescence intensity of Fluo4 (20 mins post-infection; MOI 2). **(H)** Quantification of LAMP1 recruitment to phagosomes containing *C. albicans* in IRE1 WT or IRE1^ΔR^ iBMDM with or without treatment of BAPTA-AM, a cell-permeable calcium chelator, as measured by LAMP1 mean fluorescence intensity associated with *C. albicans*-expressed iRFP. Values are the mean ± SEM from 2-4 biological replicates, as indicated by data points. *p < 0.05, **p < 0.01, *** p < 0.001, by unpaired Student’s t-test (B-D, F, G) or one-way ANOVA with Tukey’s multiple comparisons test (H).

**Supplemental Figure 2:**
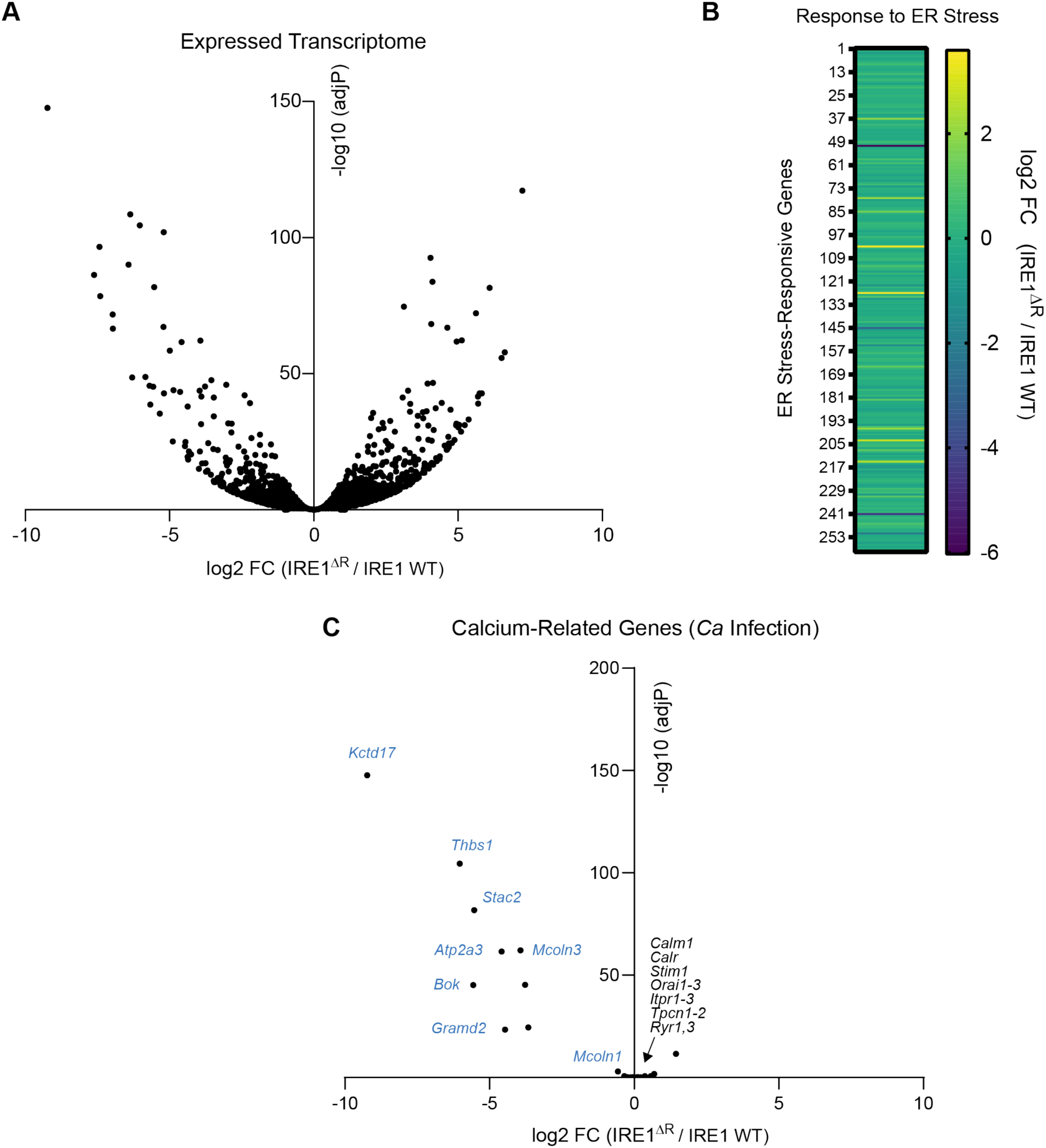
IRE1α regulates macrophage gene expression. **(A)** Volcano plot of the effect of IRE1α activity ablation on the expressed transcriptome (IRE1^ΔR^ / IRE1 WT) of iBMDM, revealed by RNA-seq. **(B)** Heatmap of differential gene expression in IRE1^ΔR^ iBMDM of genes in the GO category “Response to ER stress”, showing that IRE1^ΔR^ macrophages do not have a chronic ER stress signature. **(C)** Heatmap of downregulated genes involved in organelle calcium homeostasis (blue text) in IRE1^ΔR^ iBMDM, and major calcium homeostasis regulators (black text) whose expression is not impacted.

**Supplemental Figure 3: Related to Figure 4.**
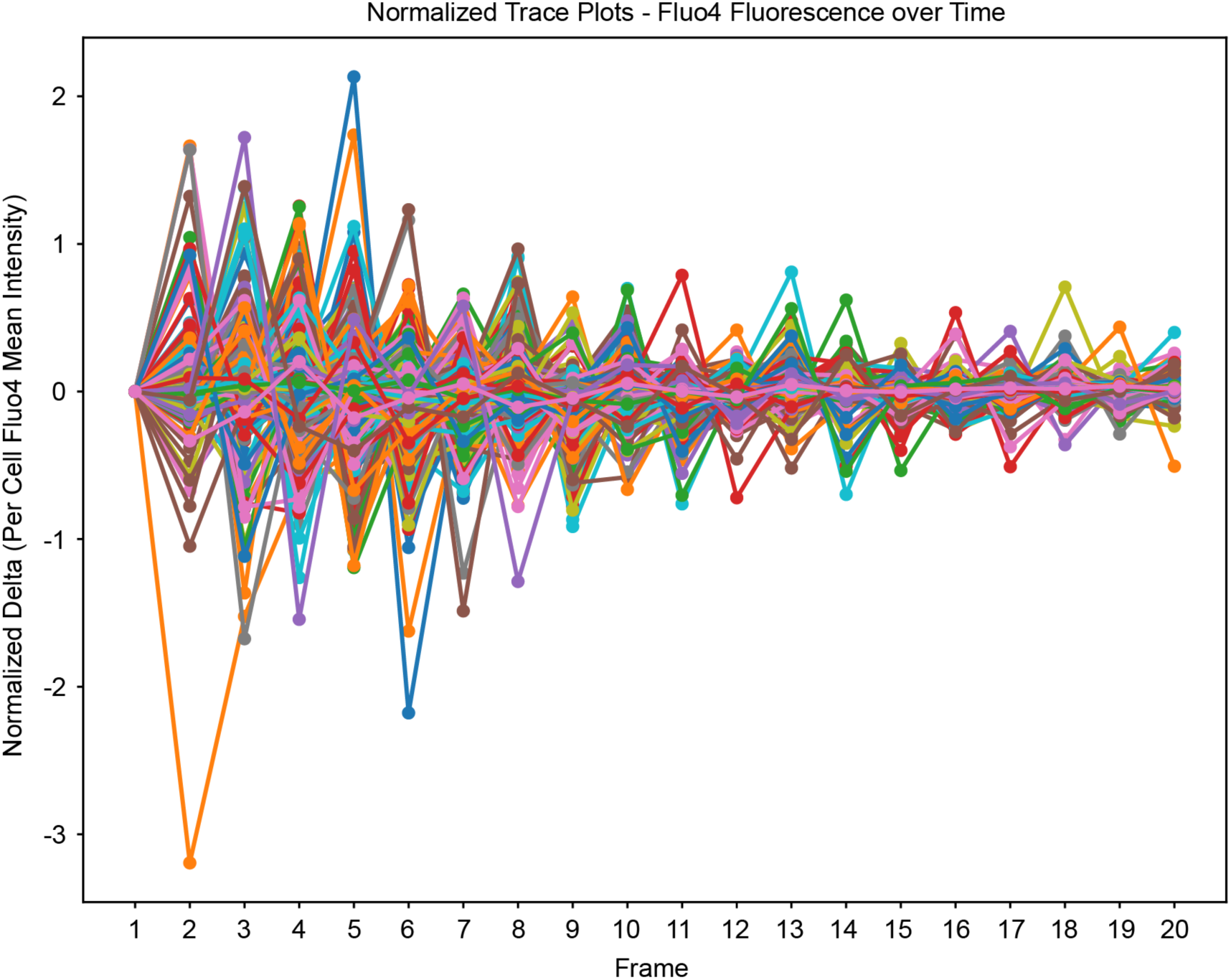
Representative trace plots of cellular Fluo4 intensity in individual cells tracked over time for quantification of cellular calcium flux, as shown in Figure 4C-D.

### IRE1α promotes phagosome integrity and macrophage fungistatic activity

As lysosome recruitment maintains the integrity of the expanding phagosome during *C. albicans* infection^44^, we reasoned that *C. albicans* may escape the phagosome more readily in IRE1^ΔR^ macrophages. To test this, we used a previously-established pulse-chase assay to measure phagosome leakage in which endosomes are pre-labeled with sulforhodamine B (SRB), allowing fusion with *C. albicans* containing phagosomes and monitoring of phagosome rupture^44,69^ (Fig. 5A). Measuring SRB association with the *C. albicans*-containing phagosome over time revealed that SRB was lost from the phagosome more rapidly in IRE1^ΔR^ macrophages, supporting the hypothesis that IRE1α activity contributes to maintenance of the *C. albicans*-containing phagosome by promoting phagolysosomal fusion (Fig. 5B). As the phagolysosomal environment restricts *C. albicans* hyphal growth^70^, we also measured hyphal growth over time in WT and IRE1^ΔR^ macrophages and found that *C. albicans* hyphal growth is increased at 4 hpi in IRE1^ΔR^ macrophages (Fig. 5C), demonstrating that IRE1α activity promotes the fungistatic activity of macrophages. Phagosome rupture during *C. albicans* infection has been associated with macrophage proinflammatory cytokine production^37,40,44,71^. Therefore, we tested secretion of IL-1β, TNF, and IL-6 from WT and IRE1^ΔR^ macrophages after LPS priming and *C. albicans* infection (Fig. 5D-F). Consistent with increased phagosome rupture observed in IRE1^ΔR^ macrophages, we also saw increased supernatant IL-1β and TNF levels, while IL-6 levels were unaffected (Fig. 5D-F). Together, these data suggest that IRE1α activity promotes phagosome integrity and macrophage fungistatic activity during *C. albicans* infection.

**Figure 5:**
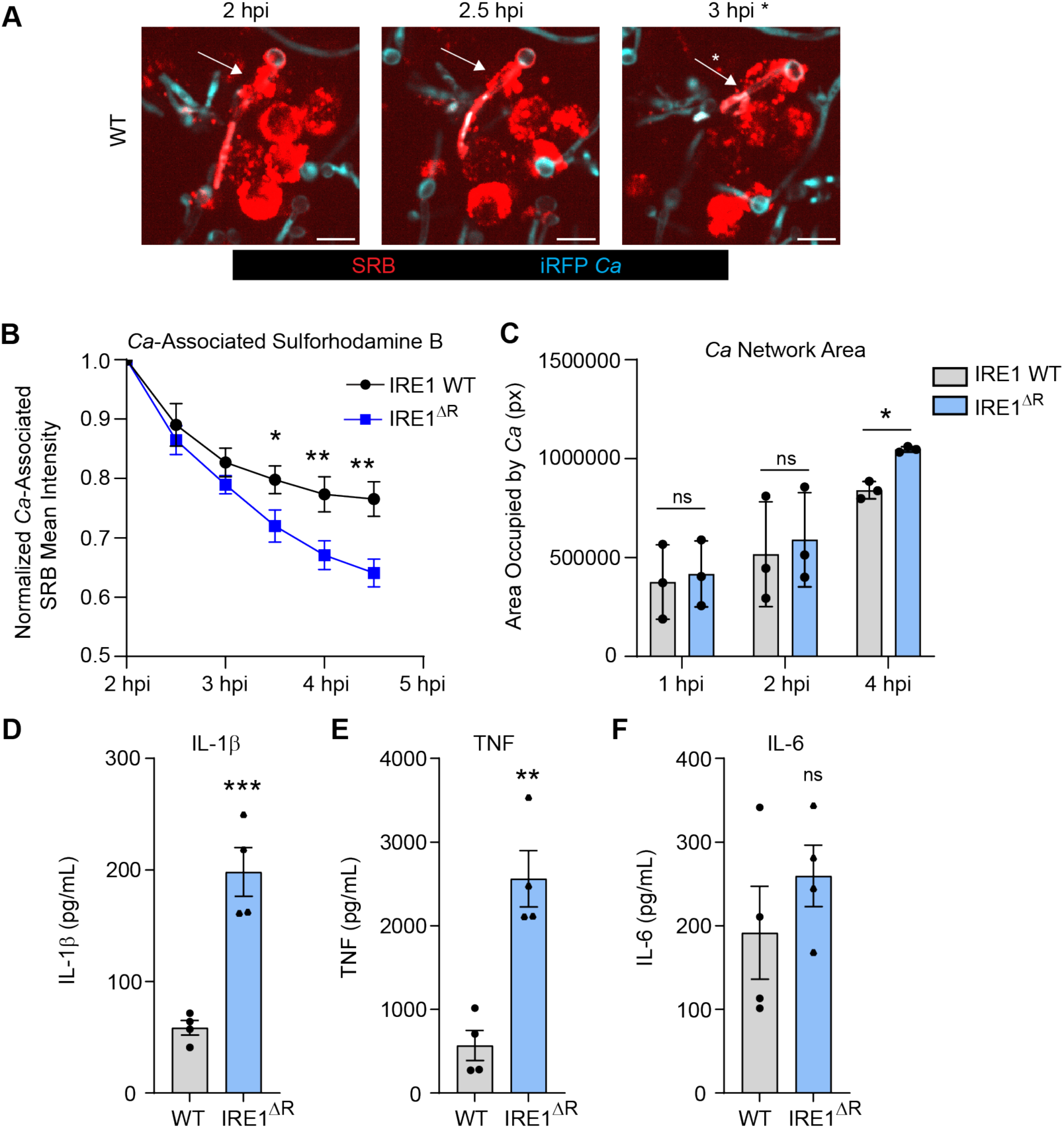
IRE1α promotes phagosome integrity and macrophage fungistatic activity. **(A)** Representative images of SRB recruitment to the phagosome containing *C. albicans*, indicated by white arrows, and loss of SRB association following phagosomal rupture at 3 hpi, indicated by white asterisk. Scale bar 10 µm. **(B)** Quantification of 3 independent experiments measuring the loss of SRB from *C. albicans* over time in IRE1 WT or IRE1^ΔR^ iBMDM. **(C)** Quantification of the area occupied (pixels) by *C. albicans* hyphae at 1, 2, and 4 hpi in IRE1 WT or IRE1^ΔR^ iBMDM. **(D-F)** Expression of proinflammatory cytokines (IL-1β (D), TNF (E), and IL-6 (F)) were measured by ELISA following 3 h of LPS priming and 5 h of *C. albicans* infection (MOI=1) in IRE1 WT and IRE1^ΔR^ iBMDM. Graphs show the mean ± SEM of 3-4 biological replicates. *p < 0.05, **p < 0.01, *** p < 0.001 by unpaired Student’s t test (B, D-F), or one-way ANOVA with Tukey’s multiple comparisons test (C).

### IRE1α promotes macrophage fungicidal activity

Escape from the phagosome likely allows *C. albicans* to evade fungicidal effectors in addition to allowing for more rapid growth. To determine whether IRE1α contributes to macrophage fungicidal activity, we measured the ability of IRE1α WT and IRE1^ΔR^ macrophages to kill phagocytosed *C. albicans*, using a dual fluorescence assay in which iRFP-expressing *C. albicans* is pre-labeled with calcofluor white (CFW) prior to macrophage infection (Fig. 6A). Live *C. albicans* express iRFP and are CFW labeled (iRFP^+^ CFW^+^), while killed *C. albicans* lose iRFP fluorescence but can be identified by CFW labeling (iRFP^-^ CFW^+^). Using this assay, we found that IRE1^ΔR^ macrophages were defective at killing phagocytosed *C. albicans*, demonstrating that IRE1α activity contributes to the fungicidal activity of macrophages (Fig. 6B).

**Figure 6:**
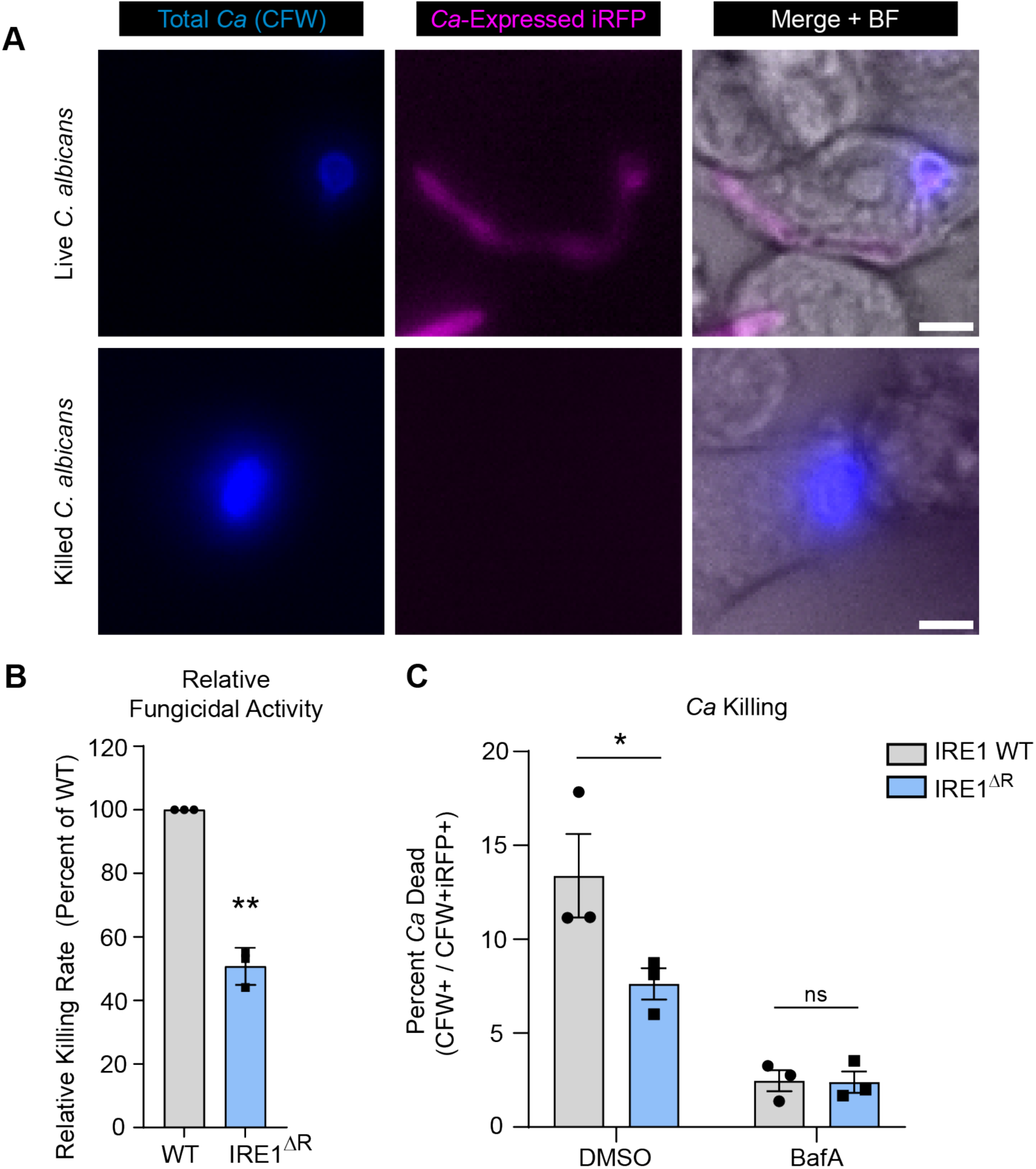
IRE1α promotes macrophage fungicidal activity. **(A)** Representative micrographs of live intracellular *C. albicans* and killed intracellular *C. albicans* within IRE1 WT iBMDM at 7 hpi. Endogenous expression of iRFP by *C. albicans* indicates viability (CFW+ iRFP+); loss of iRFP expression indicates killed *C. albicans* (CFW+ iRFP-). Scale bar 5 µm. **(B)** Quantification of 3 independent *C. albicans* killing experiments (CFW+ iRFP- / CFW+ iRFP+), relative to WT. **(C)** Graphs show the mean ± SEM of 3 biological replicates. *p < 0.05, **p < 0.01, by unpaired Student’s t test (B), or two-way ANOVA with Tukey’s multiple comparisons test (C).

To determine whether failure to recruit lysosomes to the phagosome is responsible for the fungicidal defect observed in IRE1^ΔR^ macrophages, we tested the effect of Bafilomycin A (BafA), which inhibits vacuolar ATPase activity and thus phagosome-lysosome fusion, on the ability of IRE1 WT and IRE1^ΔR^ macrophages to kill *C. albicans* (Fig. 6C). BafA treatment suppressed the fungicidal activity of both IRE1 WT and IRE1^ΔR^ macrophages, reinforcing the importance of phagolysosomal fusion for killing of *C. albicans*. Additionally, BafA treatment ablated the difference between IRE1 WT and IRE1^ΔR^ macrophages in fungicidal capacity, demonstrating that defective phagolysosomal fusion in IRE1^ΔR^ macrophages is responsible for their compromised fungicidal activity (Fig. 6C). Together, these data support a model in which IRE1α activity supports phagolysosomal fusion during *C. albicans* infection of macrophages to maintain phagosome integrity and allow killing of ingested *C. albicans*.

### IRE1α regulates cytokine production and phagocyte fungicidal activity *in vivo*

Our *in vitro* assays allowed in-depth interrogation of the role of IRE1α in interactions between *C. albicans* and bone marrow-derived macrophages. To examine the impact of IRE1α during systemic *C. albicans* infection *in vivo*, we utilized LysM-Cre to delete IRE1α activity in macrophages and neutrophils in mice (IRE1^fl/fl^ LysM^Cre^), followed by systemic infection with *C. albicans* expressing iRFP for 24 hours. Previous work demonstrated that the IRE1α-XBP1S axis in neutrophils drives fatal immunopathology starting at 5 days post-systemic *C. albicans* infection^14^. However, we found that female IRE1^fl/fl^ LysM^Cre^ mice had higher levels of serum cytokines such as IL-1Ra, TNF, and IL-6 than littermate controls (IRE1^fl/fl^) at 24 hpi (Fig. 7A). These data are in agreement with the suppressive effect of IRE1α activity on cytokine production in macrophages observed in our *in vitro* assays (Fig. 5A). Interestingly, these increased cytokine levels were observed specifically in female mice, as male mice exhibited generally similar cytokine levels to littermate controls (Fig. 7B), suggesting sex-specific roles for IRE1α during *C. albicans* infection. Additionally, we determined whether IRE1α supports the fungicidal activity of phagocytes *in vivo* using an immunofluorescence assay with dissociated kidney samples from IRE1^fl/fl^ LysM^Cre^ mice compared to IRE1^fl/fl^ controls. For this assay, *C. albicans* viability in myeloid cells was measured using an anti-*Candida* antibody to identify total *C. albicans* and iRFP to indicate viability, as well as anti-CD11b to identify leukocytes that had phagocytosed *C. albicans* within the kidney (Fig. 7C). Quantification of *C. albicans* viability within the kidney tissue revealed that while overall *C. albicans* viability in the kidney tissue was not different between IRE1^fl/fl^ LysM^Cre^ mice and IRE1 ^fl/fl^ control mice (Fig. 7D), *C. albicans* killing by phagocytic cells was less effective in mice lacking IRE1α activity (Fig. 7E), and this difference in killing efficacy between IRE1^fl/fl^ and IRE1^fl/fl^ LysM^Cre^ phagocytes appeared to be exacerbated in female mice (Fig. 7F). These data suggest that IRE1α supports the fungicidal activity of phagocytic cells *in vivo*, in agreement with our *in vitro* data, and interestingly suggest a sex-specific role for IRE1α in coordinating cytokine responses and controlling fungal infection in female mice. Together, these data establish a role for IRE1α in suppressing serum cytokine production and the fungicidal activity of phagocytes *in vivo*.

**Figure 7:**
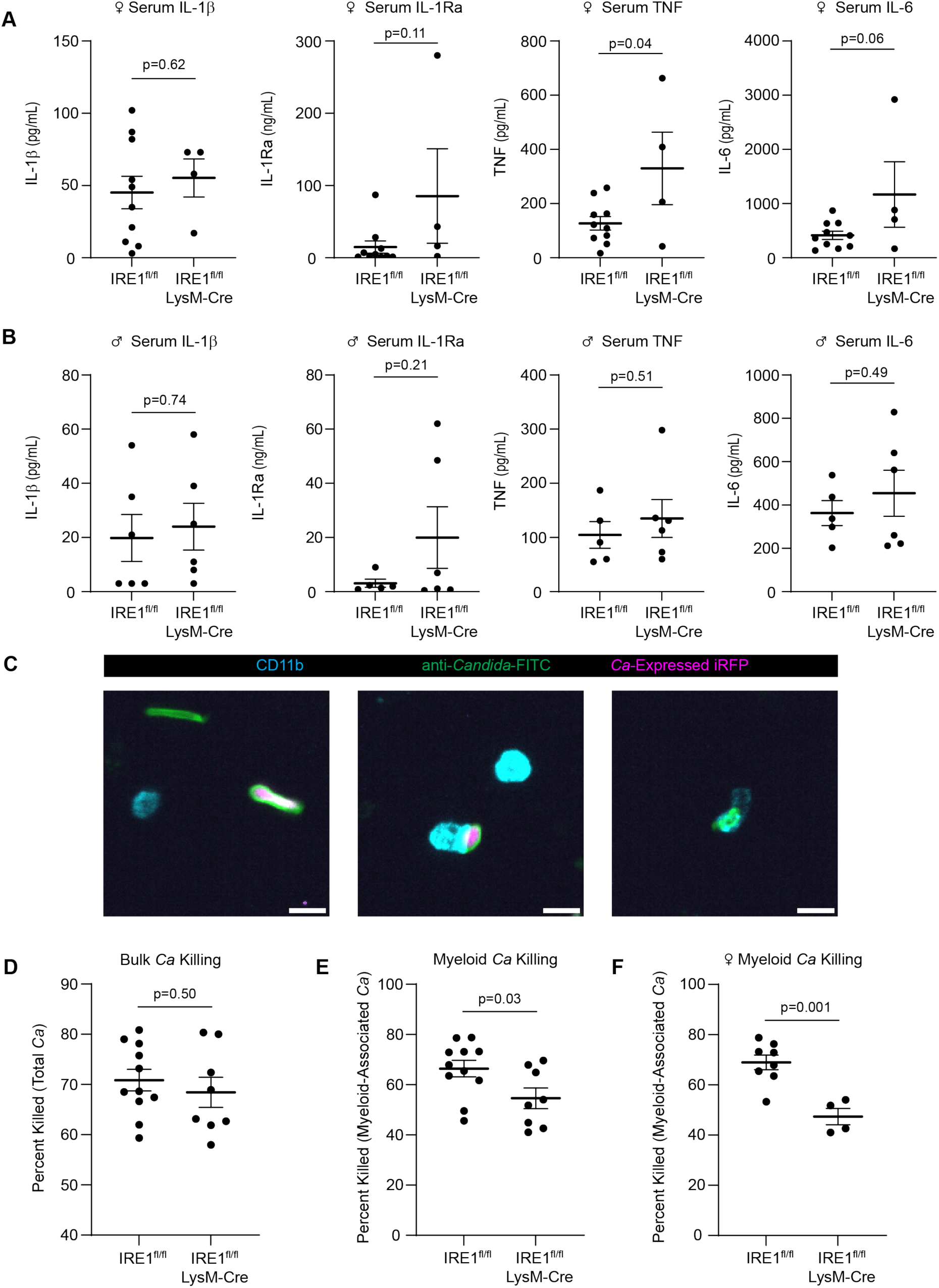
IRE1α activity in myeloid cells regulates cytokine levels and phagocyte fungicidal activity *in vivo*. **(A-B)** ELISA data from mouse serum 24 hours post-systemic *C. albicans* infection in female (A) or male (B) mice. Mice were intravenously infected with 10^6^ CFU and serum was collected through cardiac puncture. **(C)** Representative micrographs of *C. albicans* in dissociated kidney cells showing total *C. albicans* (anti-*Candida*-FITC+) in green, live *C. albicans* (anti-*Candida*-FITC+ *C. albicans*-expressed iRFP+), and CD11b positive cells to identify host leukocytes. Images show non-myeloid-associated live and dead *C. albicans* (left), myeloid-associated live *C. albicans* (middle), and myeloid-associated killed *C. albicans* (right). Scale bar 10 µm. **(D-F)** The percent of *C. albicans* killed was quantified in the kidney tissue (D), in myeloid cells from male and female mice (E), or in myeloid cells from female mice only (F). Graphs show the mean±SEM of data from individual mice. P values determined by unpaired Student’s t test.

## Discussion

Cell and organelle stress responses are crucial regulators of innate immunity and infection outcomes during bacterial and viral infection^1,2^, however, the role of these stress responses in antifungal innate immunity have not been explored in depth. Here, we show that *C. albicans* infection of macrophages results in activation of the IRE1α branch of the mammalian ER stress response. This activation is not dependent on misfolded protein stress, but instead requires signaling through the CLR pathway. Interestingly, macrophages lacking IRE1α activity have impaired fungicidal activity due to inefficient lysosome recruitment to the phagosome containing *C. albicans*, allowing phagosomal escape by *C. albicans* and likely evasion of fungicidal effectors. Together, these results demonstrate roles for IRE1α in the antifungal responses of macrophages.

IRE1α is known to be activated by bacterial and viral infections, but the mechanisms driving its activation are incompletely understood. While it has been suggested that innate immune signaling may trigger IRE1α activation independently of misfolded protein stress, this hypothesis had not been thoroughly tested. Here, we determined that CLR signaling through CARD9 triggers IRE1α activation during *C. albicans* infection. Notably, measurable protein misfolding did not precede IRE1α activation in response to either *C. albicans* infection or LPS treatment, suggesting a potential protein misfolding-independent shared mechanism of IRE1α activation downstream of innate immune signaling. However, neither TLR signaling nor TRAF6 activity were required for IRE1α activation during *C. albicans* infection, demonstrating that TLR and CLR signaling activate IRE1α through distinct mechanisms and that *C. albicans* triggers non-canonical activation of IRE1α.

A potential route of IRE1α activation in the absence of misfolded protein accumulation is post-translational modification, such as ubiquitination or phosphorylation ^21,72^. Ubiquitination of IRE1α by E3 ubiquitin ligases such as TRAF6 and CHIP contribute to IRE1α activation in response to LPS treatment or geldanamycin-induced ER stress, respectively^21,73^. However, we found IRE1α activation is TRAF6-independent during *C. albicans* infection, suggesting an alternative mechanism of activation. Additionally, the endonuclease activity of IRE1α depends on its phosphorylation status, which is governed by its own kinase activity, or in certain contexts may be triggered by other kinases^72^. We observed that CLR-mediated IRE1α activation required CARD9 in response to *C. albicans*. CARD9 forms a complex with BCL10 and MALT1, resulting in a filamentous scaffold for the assembly and activation of additional post-translational modifiers, such as the kinase TAK1 and the E3 ubiquitin ligase TRAF2, which interacts with IRE1α but is thought to act downstream of IRE1α^74^. Therefore, CARD9 activation could facilitate interaction of IRE1α with a post-translational modifier to enable its activation. As CARD family proteins and IRE1α have broad and overlapping functions in innate immune activation and immune cell function^75,76^, future work interrogating the molecular mechanisms by which CARD9 triggers IRE1α activation will be of interest.

The convergence of TLR and CLR signaling on IRE1α activation in macrophages, seemingly prior to accumulation of misfolded proteins, leads us to propose a model in which innate immune signaling induces anticipatory activation of IRE1α. This may serve to increase the secretory activity of macrophages prior to protein misfolding for efficient innate immune responses. IRE1α activity is crucial for the maturation and function of highly secretory cell types, such as plasma cells^77,78^, pancreatic beta cells^79^, macrophages^6,80^, and T cells^81^. Additionally, considering the broad roles of IRE1α and XBP1S in innate immunity, including cytokine induction^6,10^, metabolic plasticity^24^, ROS production, and microbicidal activity^12,13^, proactive strategies for IRE1α activation prior to protein misfolding may be important for innate immune regulation. Further elucidation of the mechanisms driving protein misfolding-independent IRE1α activation will identify targets for tuning of IRE1α activity, with potential for therapeutic utility for diseases in which aberrant IRE1α activity may contribute to disease progression, such as cancer and obesity^82,83^. Exploration of other pathways that may trigger non-canonical activation of IRE1α will help shape our understanding of IRE1α regulation.

When investigating the consequences of IRE1α activity during antifungal responses, we found a novel regulatory function of IRE1α in promoting transient phagosomal calcium flux. Using live imaging, we reveal transient phagosomal calcium influx during *C. albicans* infection that occurs several minutes after phagocytosis and is promoted by IRE1α. Interestingly, as IRE1α-dependent phagosomal calcium flux was observed within minutes of infection, this phenotype may reflect basal functions of IRE1α activity, rather than CLR-mediated IRE1α activation. While it has been previously shown that calcium flux is required for phagosome maturation during *C. albicans* infection^44^, and Candidalysin has been shown to induce cellular calcium flux in epithelial cells^84^, the transient accumulation of calcium in the *C. albicans* phagosome has not been described previously. Moreover, the source of this calcium and the mechanism of its uptake into the phagosome are not yet defined. Previous work has shown that IRE1α can be recruited to pathogen-containing autophagosomes^85^, and ER-phagosome contact sites can regulate phagolysosomal fusion through calcium signaling^86^. Interestingly, IRE1α can function as a scaffold at ER-mitochondria contact sites, allowing mitochondrial calcium uptake and regulation of cellular metabolism^26^. We observed that transcripts encoding genes involved in endocytosis and calcium signaling were enriched among downregulated genes in IRE1^ΔR^ macrophages (Table S1.2), highlighting the possibility that gene expression regulation by IRE1α may influence phagolysosome fusion. Whether IRE1α has broad roles in phagosome-lysosome fusion, ER-phagosome contact sites, or the degradative capacity of macrophages and other cell types will be an important future direction.

IRE1α in the myeloid compartment was shown to drive immunopathology during systemic *C. albicans* infection, and IRE1α ablation in neutrophils prolonged survival of infected hosts in a murine systemic *C. albicans* infection model^14^. In this study, it was revealed that ROS production in *C. albicans* infected neutrophils triggers protein misfolding and IRE1α activation, and subsequent XBP1S production enhanced the production of proinflammatory cytokines, driving fatal kidney immunopathology^14^. Our work complements these findings, uncovering new functions of IRE1α in macrophage antifungal responses during early stages of infection. We similarly found that stimulation of the CLR pathway can trigger IRE1α activation, although we report that protein misfolding is not required for early IRE1α activation in macrophages. Additionally, while IRE1α was not required for neutrophil fungicidal activity, we found that it does support macrophage fungicidal activity. Neutrophils are more effective at killing *C. albicans* than macrophages^87^, and presumably utilize distinct fungicidal effectors, although little is known about the mechanisms by which macrophages kill *C. albicans*^43^. A full understanding of the mechanisms by which IRE1α augments macrophage fungicidal activity will require characterization of the effectors of fungal killing in macrophages.

In summary, our work suggests that innate immune signaling can trigger non-canonical activation of IRE1α and highlights new roles for IRE1α in the fungicidal capacity of macrophages. These findings will help shape our understanding of the activation and function of IRE1α in infection and other settings. Protein misfolding-independent activation of IRE1α suggests new paradigms to explore in other contexts, such as sterile inflammation and obesity^83^, and a critical role for IRE1α as a sensor of other agents that may perturb cellular homeostasis. Dissection of the molecular mechanisms regulating early IRE1α activation may identify new therapeutic targets for regulation of IRE1α activity. Further, a better understanding of the role of IRE1α in the endocytic pathway will provide fundamental understanding of communication between the ER and endocytic compartments. Overall, these findings reveal exciting roles for a critical component of the UPR during fungal infection with broad potential impacts for our understanding of cell biology.

## Supplemental tables

Table S1: RNA-seq data from IRE1 WT and IRE1^ΔR^ macrophages infected with *C. albicans*.

- **Table S1.1:** Differential gene expression in *C. albicans* infected IRE1^ΔR^ macrophages compared to IRE1 WT
- **Table S1.2:** Gene ontology analysis of pathways enriched among genes downregulated in *C. albicans* infected IRE1^ΔR^ macrophages compared to IRE1 WT

## Supplemental Movies

**Supplemental Movie 1:** Live imaging of Fluo4 in *C. albicans*-infected IRE1 WT macrophages

**Supplemental Movie 2:** Live imaging of Fluo4 in *C. albicans*-infected IRE1^ΔR^ macrophages

## Acknowledgements

We thank the following colleagues who provided cell lines and reagents: Dr. Stacy Horner, Dr. Ling Qi, Dr. Stu Levitz, Dr. Tod Merkel, and Dr. Scott Soleimanpour, as well as Dr. Jonathan Sexton for microscopy equipment and advice, and Dr. Basel Abuaita for immortalization of IRE1^fl/fl^ ^exon20–21^ and TLR2/4/9 knockout macrophages and matched controls. We thank members of the O’Meara and O’Riordan labs for discussion. This work was supported by National Institute of Health grants K22 AI137299 (T.R.O.), R01 AI157384 (M.X.D.O.), T32AR007197 (M.J.M.), and T32AI007528 (M.J.M.). Other funding sources include the University of Michigan Pioneer Fellows Program (M.J.M.).

## Author contributions

Conceptualization: M.J.M., M.X.D.O., and T.R.O. Investigation: M.J.M., M.B.R., B.C.M., E.B.O., F.M.A., and T.L.S. Formal analysis: M.J.M., T.R.O., M.X.D.O. Software: M.J.M., M.B.R., B.C.M, E.B.O., and T.R.O. Writing – original draft: M.J.M., T.R.O., and M.X.D.O. Writing – review and editing: M.J.M., M.B.R., B.C.M., E.B.O., F.M.A., T.L.S., M.X.D.O., and T.R.O. Funding acquisition: M.J.M., M.X.D.O., and T.R.O.

## Methods

### Plasmids

pLEX-FLAG-Cre-GFP was generated by cloning PCR-amplified N-terminal FLAG tagged Cre-GFP (from pCAG-Cre-GFP; Addgene #13776) (Forward primer: TAAAGCGGCCGCTATGGCCAATTTACTGACCG; Reverse primer: CTCTAGACTCGAGTTAACTTACTTGTACAGCTCGTCCA) coding sequence into the pLEX expression vector using NotI and XhoI restriction sites. pLEX-FLAG-GFP vector was a gift from Dr. Stacy Horner. All plasmids were verified by whole plasmid sequencing (Plasmidsaurus).

### Cell lines

All cell lines were incubated at 37⁰C with 5% CO2. Bone marrow-derived macrophages (BMDM) were grown in bone marrow media (BMM), containing modification of Eagle’s medium (DMEM; Thermo Fisher Scientific) supplemented with 20% fetal bovine serum (Thermo Fisher Scientific), 30% L929 conditioned media, and 1 mM sodium pyruvate (Thermo Fisher Scientific). BMDM were immortalized (iBMDM) using J2 retrovirus^88^. L-929 cells were cultured in minimum essential Eagle’s medium supplemented with 2 mM l-glutamine, 1 mM sodium pyruvate, 1 mM nonessential amino acid, 10 mM HEPES, and 10% FBS. All experiments were performed in experimental media (RPMI supplemented with 3% FBS) unless otherwise indicated. All cell lines were verified as mycoplasma free using the Lookout Mycoplasma PCR Detection Kit (Sigma-Aldrich) and genetic identities were validated by PCR, western blotting, or functional assays. IRE1^fl/fl^ ^exon20-21^ (-/+ Cre) mice were a gift from Dr. Ling Qi, CARD9 knockout mice and littermate WT mice were a gift from Dr. Stu Levitz, TLR2/4/9 knockout mice and littermate WT mice were a gift from Dr. Tod Merkel, and TRAF6^fl/fl^ mice were a gift from Dr. Scott Soleimanpour. To generate IRE1^ΔR^ and control IRE1 WT macrophages, iBMDM from IRE1^fl/fl^ ^exon20-21^ mice with and without inducible Cre expression were treated with 4-hydroxy tamoxifen for 24 hours, followed by clonal expansion of cell lines. IRE1^ΔR^ macrophages were confirmed by immunoblotting and *Xbp1* splicing assays. To generate TRAF6 KO cell lines, first lentiviral particles encoding GFP or CRE-GFP were generated by harvesting supernatant 72 h post-transfection of 293T cells with pLEX-FLAG-GFP, or pLEX-FLAG-Cre-GFP, and the packaging plasmids psPAX2 and pMD2.G (provided by Dr. Stacy Horner). These supernatants were then used to transduce TRAF6^fl/fl^ iBMDM for 24 hours. Following transduction, cells were selected in 3 µg/mL puromycin (Sigma) for 48 hours and single cell colonies were isolated. TRAF6 deletion in CRE-GFP cell lines was verified by immunoblotting in CRE-GFP expressing TRAF6^fl/fl^ iBMDM clonal cell lines (KO-1 and KO-2). GFP-expressing TRAF6^fl/fl^ iBMDM clonal cell lines were used as a control (WT-1 and WT-2).

### *Candida albicans* infection and LPS treatment

*C. albicans* cells were cultured at 30 °C in YPD liquid media (1% yeast extract, 2% peptone, 2% dextrose) with constant agitation. All strains were maintained as frozen stocks of 25% glycerol at -80 °C. For infection of iBMDM, macrophages were seeded in experimental plates overnight at approximately 80% confluence. Experimental media (RPMI (Gibco), supplemented with 3% FBS) was inoculated with log-phase *C. albicans* cells counted for a calculated MOI of 1. LPS from *E. coli* O111:B4 (Sigma-Aldrich L2630) was diluted to 100 ng/mL in experimental media for all experiments.

### RT-qPCR

Total cellular RNA was extracted from all samples using TRIzol (Thermo Fisher Scientific), according to manufacturer’s protocol. RNA was then reverse transcribed using the iScript cDNA synthesis kit (Bio-Rad) as per the manufacturer’s instructions. The resulting cDNA was diluted 1:5 in nuclease-free H2O. RT-qPCR was performed in triplicate using the PowerUP SYBR Green PCR master mix (Thermo Fisher Scientific) and the Bio-Rad CFX Opus 384 Real-Time RT-PCR systems. *Xbp1-S* transcript was amplified using primers Forward: GCTGAGTCCGCAGCAGGT and Reverse: CAGGGTCCAACTTGTCCAGAAT. *Gapdh* transcript was amplified using primers Forward: CATCACTGCCACCCAGAAGACTG and Reverse: ATGCCAGTGAGCTTCCCGTTCAG.

### Semi-quantitative *Xbp1* splicing gel analysis

Total cellular RNA was extracted using TRIzol (Thermo Fisher Scientific), according to the manufacturer’s protocol. RNA was then reverse transcribed using the iScript cDNA synthesis kit (Bio-Rad) as per the manufacturer’s instructions. The resulting cDNA was diluted 1:5 in nuclease-free H2O. *Xbp1* transcript was amplified by PCR using primers XF and XR, followed by PCR cleanup using the Qiagen PCR Cleanup Kit. The amplified *Xbp1* product was then digested using PstI, which recognizes a cleavage site within the 26 base pair intron that is removed by IRE1α activity^89^. Following digestion, *Xbp1* bands were resolved on a 2% agarose gel and visualized by ethidium bromide staining and imaging on a BioRad gel dock.

### Immunoblotting

Cells were lysed in a modified radioimmunoprecipitation assay (RIPA) buffer (10 mM Tris [pH 7.5], 150 mM NaCl, 0.5% sodium deoxycholate, and 1% Triton X-100) supplemented with protease and phosphatase inhibitor cocktail (Millipore-Sigma) and clarified lysates were harvested by centrifugation. Quantified protein (between 5 and 15 mg) was added to a 4X SDS protein sample buffer (40% glycerol, 240 mM Tris-HCl [pH 6.8], 8% SDS, 0.04% bromophenol blue, 5% beta-mercaptoethanol), resolved by SDS/PAGE, and transferred to nitrocellulose membranes in a 25 mM Tris-192 mM glycine-0.01% SDS buffer. Membranes were stained with Revert 700 total protein stain (LI-COR Biosciences), then blocked in 3% bovine serum albumin. Membranes were incubated with primary antibodies for 2 hours at room temperature or overnight at 4C. After washing with PBS-T buffer (1 3 PBS, 0.05% Tween 20), membranes were incubated with species-specific IRDye-conjugated antibodies (Licor, 1:5000) for 1 hour at room temperature, followed by imaging on an Odyssey imaging system (LI-COR Biosciences). The following antibodies were used for immunoblotting: rabbit anti-IRE1α (CellSignaling 3924, 1:1000); rabbit anti-XBP1 (Abcam AB-37152, 1:1000); mouse anti-ACTIN (ThermoFisher ACTN05 (C4), 1:5000); rabbit anti-TRAF6 (Abcam ab40675, 1:1000); rabbit anti-CARD9 (CellSignaling 12283, 1:1000).

### Quantification of immunoblots

Following imaging using the LI-COR Odyssey imager, immunoblots were quantified using ImageStudio Lite software, and raw values were normalized to total protein (Revert 700 total protein stain) or ACTIN for each condition.

### Thioflavin T assay

iBMDM (2*10^5^ cells/well) were seeded in a 24-well plate overnight and then infected with *C. albicans*, or treated with LPS or thapsigargin for indicated timepoints. Thioflavin T (Cayman Chemical, 5 µM) was added 2 hours prior to endpoint. Cells were scraped into ice cold PBS and thioflavin T intensity was measured on a BD LSRFortessa X-20 flow cytometer.

### RNA-sequencing

iBMDM were seeded in 6-well plates overnight (10^6^ cells/well) then infected with *C. albicans* (MOI 1) or mock treated (4 h), then harvested in TRIzol reagent (Thermo Fisher) and RNA extraction was performed according to manufacturer protocol. Samples were then treated with Turbo DNase I (Thermo Fisher) according to manufacturer protocol and incubated at 37°C for 30 min, followed by phenol/chloroform extraction and ethanol precipitation overnight. RNA concentrations were then normalized. PolyA enrichment was performed and sequencing libraries were prepared and sequenced on an Illumina NovaSeq 6000 with 150 bp paired-end reads by Novogene.

### RNA-seq analysis

RNA-seq analysis was performed in Galaxy (usegalaxy.org). Reads were evaluated using FastQC and trimmed using cutadapt^90^, followed by quantification of transcripts from the GRCm38 mouse genome using Kallisto^91^. Differential gene expression between IRE1^ΔR^ and IRE1 WT macrophages following *C. albicans* infection (Table S1.1) was compared using DESeq2^92^. Gene ontology analysis was performed on significantly upregulated or downregulated genes in each data set using g:Profiler^93^.

### ELISA

iBMDM (3*10^4^ cells/well) were seeded in 96-well plates overnight, then primed with LPS (100 ng/mL) for 3 hours prior to *C. albicans* infection (MOI 1). Supernatants were collected at 5 hpi and submitted to the University of Michigan Cancer Center Immunology Core for quantification of secreted IL-1β, TNF, and IL-6.

### Quantification of phagocytosis of *C. albicans*

iBMDM (3*10^4^ cells/well) were seeded in a 96-well plastic-bottom imaging plate (PerkinElmer) overnight and then infected with *C. albicans* (MOI 1) for 30 minutes. Wells were then fixed in 4% parafolmaldehyde (Electron Microscopy Sciences) for 15 minutes, washed with PBS (ThermoFisher), and blocked with PBS containing 3% bovine serum albumin (ThermoFisher) and 5% normal goat serum (Invitrogen) for 30 minutes. FITC-conjugated anti-*Candida* antibody (LSBio LS-C103355, 1:2000) was diluted in blocking buffer and added for 1 hour with agitation to label extracellular *C. albicans*, followed by 3 5 minute washes with PBS. Wells were then permeabilized in 0.1% Triton-X 100 (Sigma-Aldrich) for 15 minutes, followed by 3 washes in PBS. Calcofluor white (Sigma-Aldrich, 1:100) was diluted in blocking buffer and added to wells for 30 minutes with agitation, followed by 3 5 minute PBS washes, and images were captured on a BioTek Lionheart FX automated microscope. A CellProfiler pipeline was developed to segment extracellular (FITC+) and total (FITC+ CFW+) *C. albicans*, and the percent phagocytosed by macrophages was calculated as 100 * (1 – (FITC+ / FITC+ CFW+)).

### Quantification of phagolysosomal fusion

iBMDM (3*10^4^ cells/well) were seeded in a 96-well plastic-bottom imaging plate (PerkinElmer) overnight and then infected with *C. albicans* SC5314 cells expressing near-infrared fluorescent protein (iRFP) driven by the *pENO1* promoter^94^ (MOI 1) at indicated timepoints. Wells were then fixed in 4% paraformaldehyde (Electron Microscopy Sciences) for 15 minutes, washed with PBS (ThermoFisher), and permeabilized in 0.1% Triton-X 100 (Sigma-Aldrich) for 15 minutes, followed by 3 washes in PBS. Wells were blocked with PBS containing 0.01% Triton-X 100, 3% bovine serum albumin (ThermoFisher) and 5% normal goat serum (Invitrogen) for 30 minutes. Primary antibodies rat anti-LAMP1 (DSHB 1D4B, 1:50) and rabbit anti-LC3 (MBL pM036, 1:400) were diluted in block buffer and added to wells for 1 hour, followed by 3 5 minute washes with PBS. Alexafluor-conjugated secondary antibodies goat anti-rat 594 and goat anti-rabbit 488 were diluted 1:500 in blocking buffer with DAPI (1:1000) and added to wells for 1 hour, followed by 3 5 minute PBS washes, and images were captured on a Yokogawa CellVoyager CQ1 automated confocal microscope. A CellProfiler pipeline was developed to segment *C. albicans* and measure the mean intensity of LAMP1 enriched at the *C. albicans* network.

### Calcium flux assay and analysis

iBMDM (3*10^4^ cells/well) were seeded in a 96-well plastic-bottom imaging plate (PerkinElmer) overnight and then loaded with Fluo-4 (1:1000; Fluo-4 Calcium Imaging Kit; Invitrogen) and CellTracker Red (1:2000; Invitrogen) according to manufacturer protocol for 20 minutes at 37⁰C, then 20 minutes at room temperature. Staining media was then removed, followed by a wash with room-temperature media. Cells were infected with *C. albicans* SC5314 cells expressing near-infrared fluorescent protein (iRFP) driven by the *pENO1* promoter^94^ (MOI 2), immediately followed by live imaging captured on a Yokogawa CellVoyager CQ1 automated confocal microscope with incubation at 37⁰C with 5% CO2. Images were captured every 90 seconds for 1 hour.

Analysis of initial Fluo-4 intensity was performed on time 0 images using a CellProfiler pipeline to identify cells and measure the mean fluorescence intensity of Fluo-4. For analysis of cellular calcium flux, the Python package spacr (https://github.com/EinarOlafsson/spacr) was used to segment and track cells over time and quantify single cell calcium oscillations. Cells were delineated with the Cellpose cyto model^95^ from CellTracker Red staining. Centroids of identified cell objects were tracked using the Trackpy particle-tracking algorithm^96^. Fluo-4 mean intensity values were normalized between 0 and 1 and corrected for photobleaching across the time series using an exponential decay model to enable the detection of calcium spikes above a threshold of 0.25 with the find_peaks function from scipy^97^. Peaks were then enumerated and characterized by collecting peak frequency and amplitude for each condition.

Analysis of phagosomal calcium influx was performed at 20 minutes post-infection using NIH Fiji/ImageJ. The line tool was used to calculate the mean fluorescence intensity of Fluo4 rings within *C. albicans*-containing phagosomes, which were measured relative to the mean fluorescence intensity of the whole parental macrophage. Calcium-high phagosomes were defined as phagosomes with Fluo-4 intensity >1.25-fold higher than the mean fluorescence intensity of the parent macrophage.

### Macrophage fungicidal activity assay

iBMDM (3*10^4^ cells/well) were seeded in a 96-well plastic-bottom imaging plate (PerkinElmer) overnight. Approximately 10^7^ *C. albicans* SC5314 cells expressing iRFP from an overnight culture were stained with calcofluor white (CFW; 100 µg/mL) for 10 minutes in the dark. Cells were then washed twice with PBS prior to macrophage infection at MOI = 1. Images of infected cultures were captured every 20 minutes on a BioTek Lionheart FX automated microscope with incubation at 37C and 5% CO2. Fungal killing was quantified at 7 hours post-infection by calculating killed *C. albicans* (iRFP^-^ CFW^+^) over total *C. albicans* (iRFP^-/+^ CFW^+^), with at least 200 *C. albicans* cells counted per condition.

### LysoSensor

iBMDM (2*10^5^ cells/well) were seeded in a 24-well plate overnight and then infected with *C. albicans* for 2 hours prior to addition of LysoSensor Yellow/Blue DND-160 (Thermo Fisher, 500 nM) for 2 minutes in experimental media. Wells were then washed 3 times in ice-cold PBS and scraped for plate reader analysis. Suspended cells were added to a black-bottom 96-well plate and absorbance and emission were measured at 329 nm Abs, 440 nm Em and 384 nm Abs, 540 nm Em to measure fluorescence intensity in high and low pH environments, respectively. The intensity of the low pH measurement was divided by the intensity of the high pH measurement, and these results were normalized to IRE1 WT Mock to determine the relative acidity of each condition.

### Sulforhodamine B Assay and *C. albicans* hyphal length measurement

iBMDM (3*10^4^ cells/well) were seeded in a 96-well plastic-bottom imaging plate (PerkinElmer) overnight, then sulforhodamine B (SRB) (Sigma-Aldrich, 150 µg/mL) was added to wells for 1 hour. SRB was then washed out and wells were with *C. albicans* expressing iRFP (MOI 1) and live imaging was performed on a Yokogawa CellVoyager CQ1 automated confocal microscope every 30 minutes for 5 hours. A CellProfiler pipeline was developed to segment *C. albicans* and measure the total area covered by hyphae, and the mean intensity of SRB enriched at the *C. albicans* network was measured at each timepoint.

### *In vivo* systemic *C. albicans* challenge experiments

Overnight cultures of *C. albicans* expressing iRFP were sub-cultured at a starting OD600 of 0.1 and grown for 4 hours, then pelleted by centrifugation and resuspended in PBS for delivery to the bloodstream of mice. 8-12 week old male and female mice lacking IRE1α activity in macrophages and neutrophils (IRE1^fl/fl^ LysM^Cre^) and littermate controls (IRE1^fl/fl^) were systemically infected with iRFP-expressing *C. albicans* (10^6^ CFU) by retro-orbital injection. At 24 hours post-infection, mice were euthanized and serum was collected by cardiac puncture, followed by isolation of serum using centrifugation of serum collection tubes. Serum samples were submitted to the University of Michigan Cancer Center Immunology Core for quantification of secreted IL-1β, IL-1Ra, TNF, and IL-6 by ELISA. Kidneys were isolated and dissociated by mechanical separation through a 70 µm cell strainer, followed by red blood cell lysis (eBioscience 10X RBC Lysis Buffer). To quantify *C. albicans* viability in kidney samples, 2*10^6^ cells per sample were subjected to immunofluorescence staining. Total *C. albicans* was stained using a FITC-conjugated anti-*Candida* antibody (1:1000; Meridian Bioscience), and myeloid cells were stained with Brilliant Violet 421-conjugated anti-CD11b antibody (1:100; Biolegend) for 1 hour in the dark with gentle agitation. After immunostaining, samples were plated in 96-well plastic-bottom imaging plates (PerkinElmer) coated with poly-D-Lysine (Gibco) and imaging was performed on a Yokogawa CellVoyager CQ1 automated confocal microscope. A CellProfiler pipeline was developed to segment *C. albicans* and host myeloid cells. Total *C. albicans* were identified from kidney tissue and myeloid cells using FITC signal, and viability was measured using iRFP intensity.

### Lead Contact and Materials Availability

Further information and requests for resources and reagents should be directed to and will be fulfilled by the Lead Contacts, Teresa O’Meara and/or Mary O’Riordan

### Data Availability

All raw data related to RNA-seq are available through GEO (accession number: GSE244303). All raw data related to microscopy are available upon request.

### Code Availability

CellProfiler pipelines for image quantification are available as Supplementary Materials (files S2-S5). Software for cellular calcium flux analysis are available from GitHub (https://github.com/EinarOlafsson/spacr).

